# Novel Bayesian Networks for Genomic Prediction of Developmental Traits in Biomass Sorghum

**DOI:** 10.1101/677179

**Authors:** Jhonathan P. R. dos Santos, Samuel B. Fernandes, Roberto Lozano, Patrick J. Brown, Edward S. Buckler, Antonio A. F. Garcia, Michael A. Gore

## Abstract

The ability to connect genetic information between traits over time allow Bayesian networks to offer a powerful probabilistic framework to construct genomic prediction models. In this study, we phenotyped a diversity panel of 869 biomass sorghum (*Sorghum bicolor* (L.) Moench] lines, which had been genotyped with 100,435 SNP markers, for plant height (PH) with biweekly measurements from 30 to 120 days after planting (DAP) and for end-of-season dry biomass yield (DBY) in four environments. We evaluated five genomic prediction models: Bayesian network (BN), Pleiotropic Bayesian network (PBN), Dynamic Bayesian network (DBN), multi-trait GBLUP (MTr-GBLUP), and multi-time GBLUP (MTi-GBLUP) models. In 5-fold cross-validation, prediction accuracies ranged from 0.48 (PBN) to 0.51 (MTr-GBLUP) for DBY and from 0.47 (DBN, DAP120) to 0.74 (MTi-GBLUP, DAP60) for PH. Forward-chaining cross-validation further improved prediction accuracies of the DBN, MTi-GBLUP and MTr-GBLUP models for PH (training slice: 30-45 DAP) by 36.4-52.4% relative to the BN and PBN models. Coincidence indices (target: biomass, secondary: PH) and a coincidence index based on lines (PH time series) showed that the ranking of lines by PH changed minimally after 45 DAP. These results suggest a two-level indirect selection method for PH at harvest (first-level target trait) and DBY (second-level target trait) could be conducted earlier in the season based on ranking of lines by PH at 45 DAP (secondary trait). With the advance of high-throughput phenotyping technologies, our proposed two-level indirect selection framework could be valuable for enhancing genetic gain per unit of time when selecting on developmental traits.

---

The development of renewable energy resources from biomass crops is a key step towards the establishment of a sustainable agroecosystem (Foley *et al.* 2011; Mace *et al.* 2013). Among the plant species amenable to bioenergy production, sorghum [*Sorghum bi-color* (L.) Moench] is a prominent candidate for genetic improvement because it is diploid (2*n*=2*x*=20), fixes carbon through the *C*_4_ pathway, predominantly autogamous, and resilient against biotic and abiotic stresses (Vermerris 2011; Lawrence and Walbot 2007). Although substantially more economic investments have been made in crops such as maize, which resulted in improvements of up to 4-fold in grain yield over the last century, proportional gains in yield might be achievable in biomass sorghum (Mullet *et al.* 2014). Currently, sorghum has an average biomass yield of 12-15 dry Mg ha^−1^ under rainfed conditions, with a predicted potential yield of 55-60 dry Mg ha^−1^ for ideotypes (Mullet *et al.* 2014). Because of its biology, sorghum has the potential to become a model organism to understand the genetic basis of growth traits related to biomass production (Brenton *et al.* 2016).

Genomic prediction (GP) is a statistical approach that predicts the unobserved phenotypes of individuals using genomic information (Meuwissen *et al.* 2001). Because of its potential to enrich for promising selection candidates, GP is increasingly becoming an important component of plant breeding and genetic resources conservation programs. In sorghum, Yu *et al.* (2016) showed that GP can optimize the management and evaluation of accessions from gene banks through the prediction of different traits. Briefly, the GP procedure involves two steps: (i) phenotyping and genotyping a reference population (training set) to train statistical models, and (ii) genotyping of unevaluated individuals (test set) for predicting their unobserved phenotypes with the trained models (Heslot *et al.* 2015). To support this procedure, plant breeding programs collect phenotypes from training set individuals evaluated in multi-environment trials. In parallel, high density single-nucleotide polymorphism (SNP) markers are scored on individuals in the training and test sets with skim sequencing or SNP arrays (Elshire *et al.* 2011; Davey *et al.* 2011; Buckler *et al.* 2016).

Mixed linear models, hierarchical Bayesian models with informative priors, kernel methods, and neural nets are modeling approaches used for GP, but minimal differences in predictive performance are typically seen across these approaches (de Los Campos *et al.* 2013; Heslot *et al.* 2015; dos Santos *et al.* 2016a). This outcome may be explained by the high density of covariates (SNP markers) compared to the population size used for training the models. This scenario is known as the large *p* and small *n* problem (*p* ≫ *n*) (Gianola *et al.* 2009). In statistical models, this may lead to the problem of multicollinearity, i.e., multiple covariates with redundant information. As it relates to GP models, markers in complete or near complete linkage disequilibrium provide redundant information and will not contribute to enhancing statistical power (Gianola 2013). Dimensionality reduction techniques, such as the artificial bins approach, may help circumvent the challenge of multicollinearity with minimal information loss, as well as mitigate the computational cost often associated with GP (Xu 2013).

The vast majority of GP studies conducted in crop species have only tested models for predicting individual traits. However, recent studies have shown the advantages of combining multiple correlated traits in a GP model (Calus and Veerkamp 2011; Jia and Jannink 2012; Fernandes *et al.* 2018), allowing genetic correlations among secondary traits to be leveraged for improving predictions of a target trait (dos Santos *et al.* 2016b; Okeke *et al.* 2017). Most of these efforts used multi-trait GBLUP—a type of multivariate mixed linear model that incorporates a genomic relationship matrix (Gianola *et al.* 2015). Despite the advances obtained so far, the use of genetic models that exploit information between traits using other parametrizations beyond those reliant on genetic correlations under multivariate normal distribution assumptions have yet to be addressed. Indeed, novel genetic models with parametrizations to partition genetic effects influencing only a single trait from those acting on multiple traits (i.e., pleiotropy) may help to better understand the genetic architecture of correlated traits.

There have been significant advances in field-based high-throughput phenotyping (HTP) technologies for the rapid measurement of plant traits over the growing season (Bao *et al.* 2019; Pauli *et al.* 2016). Measuring phenotypes at multiple time points over the life cycle of a plant can better describe the progression of growth and development (Muraya *et al.* 2017). Having collected phenotypic information on a time axis may help to identify key environmental stress events during the growing season, which might be masked if phenotypic data are only obtained at harvest (Campbell *et al.* 2018). Furthermore, the underlying genetic signals of these phenotypic responses are additional sources of information to more powerfully predict and dissect the genetic architecture of developmental plant traits (Muraya *et al.* 2017; Campbell *et al.* 2018). Statistical models that exploit temporal genetic trends are especially needed for longitudinal (repeated measure) data collected by field-based HTP systems. Such models could be used to reduce generation time and prioritize which breeding populations to evaluate.

Among the models available for analyzing traits in a time series, probabilistic graphical models (PGMs) offer a versatile, efficient, and intuitive approach for drawing inferences (Murphy 2013; Bishop 2013). Popular PGMs include directed graphical models or Bayesian networks (BNs), undirected graphical models or Markov random fields, chain models, and factor graphs (Hamelryck 2012). In particular, BNs provide the flexibility to model repeated measure and correlated trait data, as would be important for the study of developmental traits. A BN is defined as a structured graphical representation of joint distributions factored into a set of conditional probability distributions, where shaded and unshaded nodes represent known and unknown variables, respectively, and arrows showing dependence between them (Bishop 2013). The Markov condition is a key property of the BN, ensuring that a variable (child) is only dependent on the information of its parents in the network (Su *et al.* 2013; Bishop 2013). Through their ability to connect joint probability distributions, BNs enable the aggregation of advantages from multiple machine learning approaches under a directed acyclic graph structure. Notably, BNs have been diversely applied in genetic and genomic studies (Loman *et al.* 2015; Serang *et al.* 2012; Garcia *et al.* 2013; Han *et al.* 2012; Su *et al.* 2013; Neapolitan *et al.* 2013), but to our knowledge have never been used for modelling trends of genetic effects considering repeated measures and correlated traits.

There are several features of BNs that enable them to recover information from correlated data types such as multiple correlated traits scored at a single time point or the repeated measurement of a single trait across multiple time points (Bae *et al.* 2016). Several different GP models could be unified for leveraging pleiotropy or temporal genetic effects in a single BN to improve prediction accuracies. This is because these genetic effects can be modeled with a BN through connections between likelihood functions. Also, BNs offer the possibility to use general Markov chain Monte Carlo (MCMC) methods to obtain solutions for complex time series and multiple trait models that otherwise would have been mathematically intractable to derive analytically. Furthermore, the posterior samples of genomic estimated breeding values (GEBVs) may be used to create indices for understanding the uncertainty of selecting promising lines either earlier in the season or through indirect selection based on the ranking of the lines at other measurement time points or with correlated traits.

With sorghum as a model biomass crop, we developed PGMs for the GP of developmental traits in a sorghum diversity panel of nearly 900 lines. Herein, we aimed to (i) develop PGMs for the GP of plant height (PH) and dry biomass yield (DBY) traits by connecting genetic effects across multiple developmental time points and traits, and (ii) describe growth dynamics based on the change of the ranking of lines across multiple time points and correlated traits to design novel breeding strategies to genetically improve biomass sorghum.

## MATERIALS AND METHODS

### Plant material, field experiments and phenotypic data

In this study, we evaluated a biomass sorghum diversity panel consisting of 869 lines (Valluru *et al.* 2018). The diversity panel was grown at three field locations only a few km from the main campus of the University of Illinois Urbana-Champaign in 2016 (Fisher and Energy Farms) and 2017 (Maxwell and Energy Farms). Each of the four environments had one complete replication of the field experiment that contained 960 four-row plots laid out in an 40-row by 24-column arrangement. The experimental field design consisted of 16 incomplete blocks, and each block was augmented with a common set of six lines (four shared between years): Pacesetter, PI276801, PI148089, PI524948, NSL50748, and PI148084 in 2016 and Pacesetter, PI276801, PI148089, PI524948, PI525882, and PI660560 in 2017. Plots were 3 m in length, with a 1.5 m alley at the end of each plot. Plots had a spacing between rows of 0.76 m. The plant population had a targeted density of 270,368 plants ha^−1^. Experiments were planted in late May and harvested in early October. PH was measured in centimeters (cm) from the soil line to the topmost leaf whorl. A single plant was measured in each plot on a biweekly basis from 30 to 120 days after planting (DAP). Plots were harvested for above ground biomass using a four-row Kemper head attached to a John Deere 5830 tractor. Wet weight of total biomass (lbs) and biomass moisture (%) in the center two rows of each 4-row plot were measured using a plot sampler that had a near infrared sensor (model 130S, RCI engineering). DBY in dry tonne per hectare was calculated as follows: dry t ha^−1^ = total plot wet weight (kg) × (1-plot moisture) / (plot area in square meters/10,000).

### Phenotypic data analysis

Phenotypic measurements for DBY (dry t ha^−1^; one measurement at harvest) and PH (cm; one measurement on each of seven plant developmental stages) were analyzed individually with the following mixed linear model:

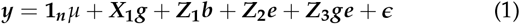

where *y* (*n* × 1) represents the phenotypic vector with *n* entries, **1**_*n*_ a unit vector, *μ* a scalar to map the population mean, *X*_**1**_(*n* × *q*) the design matrix of the *q* fixed genetic effects (number of lines), *Z*_**1**_(*n* × *l*) the design matrix of the *l* random block within environment effects, *Z*_**2**_(*n* × *s*) the design matrix of the *s* random environment (location x year combination) effects, *Z*_**3**_(*n* × *m*) the design matrix of the *m* random genotype-by-environment effects; and *g*, *b*, *e*, and *ge* are column vectors mapping the design matrices effects, respectively, and *ϵ* (*n* × 1) the vector of errors. The model random effects *b*, *e*, *ge*, and *ϵ* were assumed to follow a 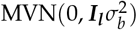, 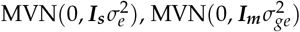, and 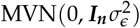, respectively.

Empirical estimates of variance components were obtained by the maximization of the restricted (residual or reduced) maximum likelihood (REML) function (Patterson and Thompson 1971). For each phenotype, the adjusted mean for each line was derived by the sum of the population mean with the fixed genetic effect of the i^th^ line 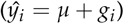.

Heritability on an line-mean basis was estimated for each phenotype. Variance component estimates were obtained by refitting model (1) with all terms as random effects in ASReml-R version 3.0 (Butler *et al.* 2009). The variance component estimates from each model for a phenotype were used to estimate heritability on a line-mean basis as the ratio of genetic variance to phenotypic variance following (Holland *et al.* 2003; Hung *et al.* 2012). Standard errors of the heritability estimates were calculated with the delta method (Lynch *et al.* 1998; Holland *et al.* 2003) in the *nadiv* R package (Wolak 2012). The Pearson’s correlation coefficient (*r*) was used to assess the degree of relationship between adjusted means for each pair of traits.

### Genotypic data

We genotyped the sorghum diversity panel using the genotyping-by-sequencing (GBS) procedure (Elshire *et al.* 2011) based on the *Pst*I-HF/*Hin*P1I and *Pst*I-HF/*B f a*I restriction enzymes. A total of 367 million sequence reads were generated (100 bp length) on a HiSeq 4000 sequencer. Sequence reads were aligned to the *Sorghum bicolor* genome v3.1 (www.phytozome.jgi.doe.gov) using Bowtie2 (Langmead and Salzberg 2012). The TASSEL 3 GBS pipeline (Glaubitz *et al.* 2014) was then used to call variants. Only biallelic SNPs were retained. Additionally, lines with >80% missing data (sample call rate) and SNPs with >60% missing data (SNP call rate) were removed. Also, SNPs with a minor allele frequency less than 5% were discarded. Missing genotypes of SNP markers were imputed using *Beagle 4.1* (Browning and Browning 2016) with default parameters and an Ne of 150,000. In total, 100,435 SNP markers were scored and converted to dosage format (0,1,2). Of the 869 total lines, 839 had both phenotypic and genotypic data; therefore, the GP analyses focused on only these 839 lines.

### Artificial bins

Due to the high dimensionality of the SNP marker matrix (100,435 loci), we developed a strategy similar to that proposed by Xu (2013) for obtaining artificial bins. In our approach, after centering (subtracting) the marker scores 2 (MM), 1 (Mm), and 0 (mm) by 2*p*, instead of averaging the columns from equally sized slices of the marker matrix, we conducted a principal component analysis (PCA) to calculate the first PC of each artificial bin. The procedure was based on the following steps: (i) subdivision of the centered marker matrix into 1000 column-slices, with each slice comprising ~100 columns; (ii) singular value decomposition of each matrix column slice into singular vectors and values; and (iii) construction of each artificial bin as the first PC coordinates of its respective matrix column slice, given by *u*_**1**_*λ*_1_, where *u*_**1**_ is the first left singular vector of the matrix slice and *λ*_1_ its first singular value. This procedure resulted in 1,000 artificial bins. This number of artificial bins was selected as a balance between model run time and predictive performance for the computationally intensive Bayesian models. In theory, the first PC (artificial bin) will retain as much information as possible from the matrix slice in one dimension in a least-square reconstruction error sense (Goodfellow *et al.* 2016).

### Probabilistic graphical models

We developed three different Bayesian models for genomic prediction of the PH and DBY traits (Figure 1). The first model, Bayesian network (BN), neither recovered information between traits nor time points. The BN has the following conditional normal likelihood form:

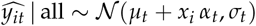

where 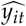 is the adjusted mean related to the i^th^ line evaluated in the t^th^ time point (or DAP), *μ*_t_ the unknown population mean in the t^th^ time point, *x*_*i*_ the known row vector (1 × *p*) of the *p* artificial bins of the i^th^ line, *α*_*t*_ the column vector (*p* × 1) of the unknown *p* artificial bins effects within in the t^th^ time point and *σ*_*t*_ the unknown residual standard deviation mapping the uncertainty around the expected value in the t^th^ time point.

**Figure 1.**
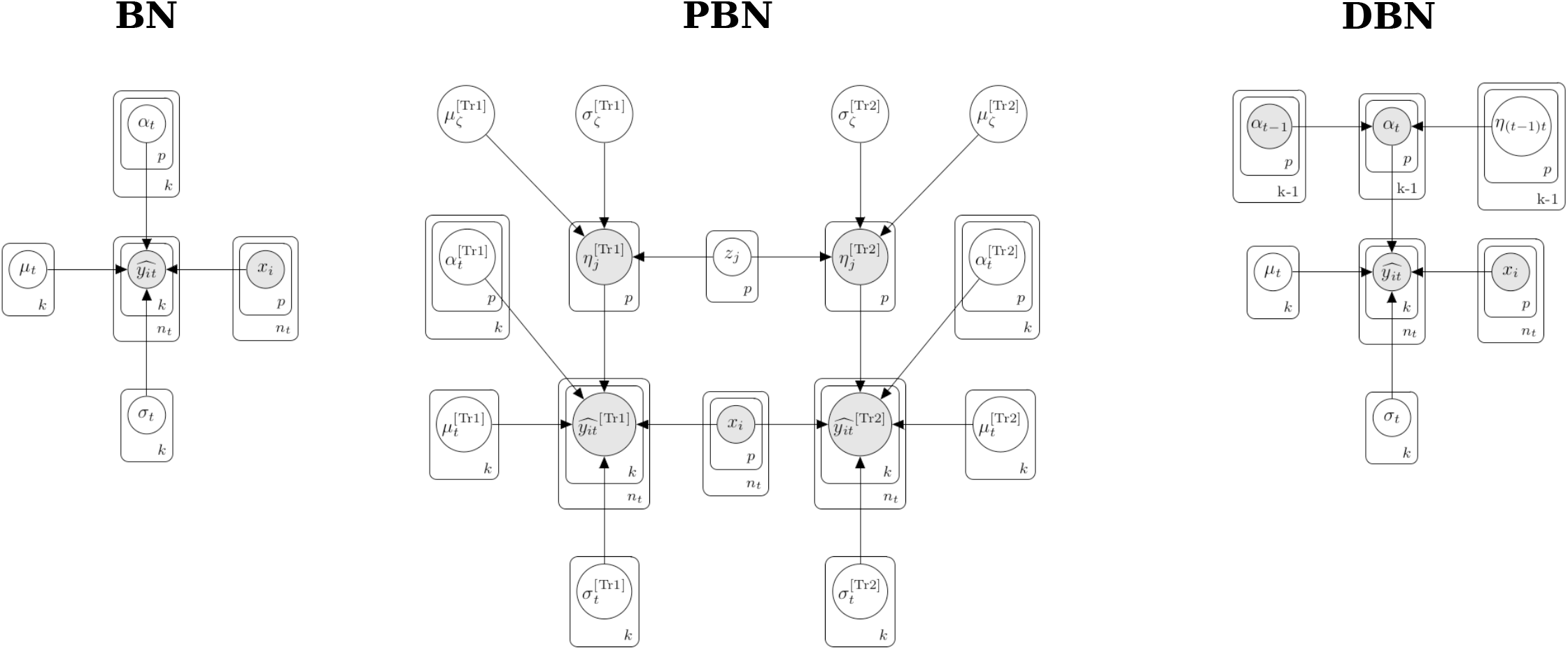
Overview of the Bayesian models. Bayesian network (BN), pleiotropic Bayesian network (PBN), and dynamic Bayesian network (DBN) probabilistic graphical models. *k*: number of time points; *n*_*t*_: number of lines within a time point; *p*: number of artificial bins; 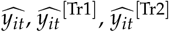: Adjusted means for the i^th^ line evaluated in the t^th^ time point, which can be for trait 1 (Tr1) or trait 2 (Tr2); *μ*t, 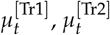: population means; *x*_*i*_ : row vector with artificial bins; *α*_*t*−1_,*α*_*t*_,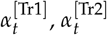: column vector with artificial bins effects; 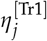, 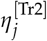: pleiotropic j^th^ bin effect; *z*_*j*_: standardized pleiotropic j^th^ bin effect; 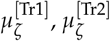: pleiotropic means hyperparameters; *η*_(*t*−1)*t*_: bin effects between the current and previous time point; 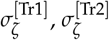 : pleiotropic standard deviations hyperparameters; *σ*_*t*_,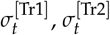 : standard deviations.

The BN has an unnormalized joint posterior distribution (hyperpriors omitted from Figure 1 for simplicity),

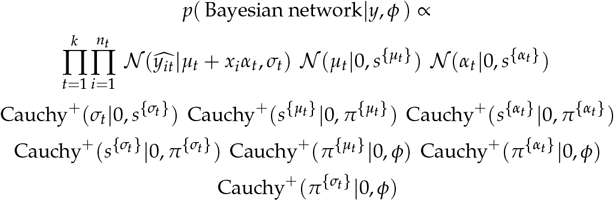

where *k* is the total number of time points, *n*_*t*_ is the total number of lines in the t^th^ time point, 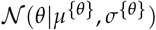 and Cauchy^+^(*θ*|*μ*^{*θ*}^, *σ*^{*θ*}^) denotes the normal probability density function, and Cauchy probability density function truncated to the real positive space 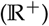 of the random variable *θ* (general notation), respectively, parametrized by the mean (*μ*_*θ*_), standard deviation (*σ*_*θ*_). The joint distribution was parameterized as second (*s*^{*θ*}^) and third (*π*^{*θ*}^) level scale hyperparameters. The known global hyperparameter was defined by *φ* = ∥*y*∥_∞_ × 10 = arg max(*y*) × 10, resulting in weakly informative second-level hyperpriors that eliminated the subjectiveness to define hyperparameters when choosing first-level prior hyperparameters (Gelman *et al.* 2014). This same approach was used for the next set of described models.

The second Bayesian model, pleiotropic Bayesian network (PBN), exploited information between PH and DBY (Figure 1). This model has two conditionally dependent normal likelihood functions that characterized the observed adjusted means distribution for each trait as follows:

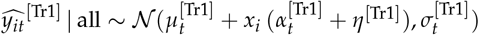

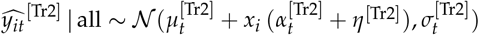

where all variables are the same from the previous model, except the column vectors *η*^[Tr1]^ (*p* × 1) and *η*^[Tr2]^ (*p* × 1), that represent the pleiotropic effects of known bins with continuous space corrected by the transformation of an unknown pleiotropic standardized random variable *z*_*j*_ for the j^th^ bin,

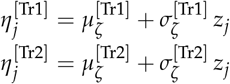

with 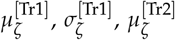, and 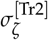 being unknown random variables. The PBN model has an unnormalized joint posterior density function (Figure 1),

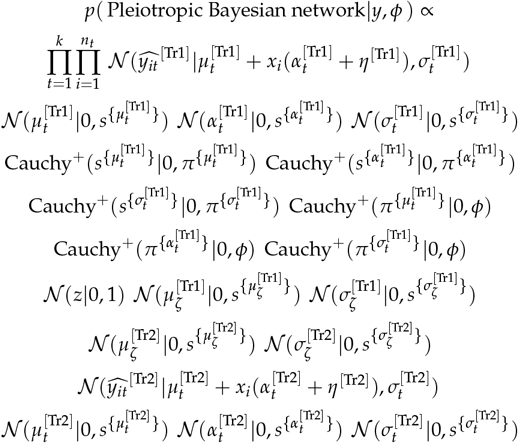

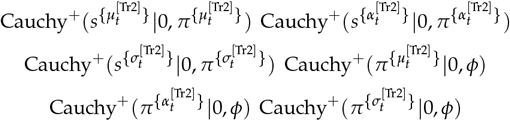

The third Bayesian model, dynamic Bayesian network (DBN), recovered information from PH measurements across multiple time points (Figure 1). This network architecture has a specific conditionally, dependent normal likelihood function for each time point as follows:

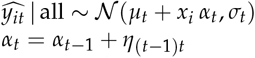

where the column vector *α*_*t*_ (*p* × 1) is the known artificial bins effects at time *t*, that are a linear combination of the *α*_*t*−1_ (*p* × 1) known artificial bins effects displayed in the previous time point (*t* − 1) plus the unknown *η*_(*t*−1)*t*_ (*p* × 1) random noise mapping the bin effect between the current and previous time points, such that genetic information is propagated over time. The artificial bins effects were treated as unknown random variables only at the first time point. The DBN model has an unnormalized joint posterior distribution (Figure 1),

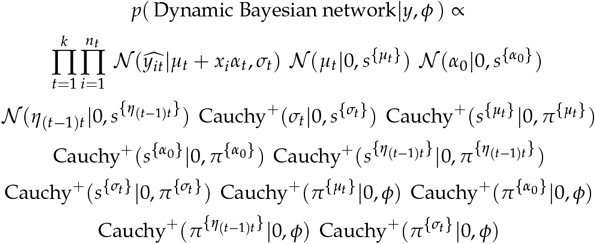

The joint distributions of the BN, PBN, and DBN models were integrated using the No-U-Term sampler algorithm available in the probabilistic programming language Stan (Hoffman and Gelman 2014). We used the implementation available in the Python package *pystan 2.17.1.0* (Team 2018). Stan compiles the probabilistic programming code in C++ and has a user interface within the Python environment. The probabilistic programming language saved time during customization of the C++ code, allowed rapid implementation during model design and training, as well as facilitated the manipulation of posterior draws after fitting the model in Python (Carpenter *et al.* 2017). We set up the No-U-Term sampler to iterate 400 times and used as warm up 50% of the samples from four Markov chains. The number of iterations (400) was selected in consideration of runtime and predictive performance.

### Multivariate GBLUP model

For comparison to the three Bayesian models, we also evaluated two different formulations of the multivariate GBLUP model (Henderson and Quaas 1976; dos Santos *et al.* 2016b; Fernandes *et al.* 2018) that recovered information between traits and/or time points. In the first formulation, only PH measurements across time points were used (MTi-GBLUP). In the second, DBY and PH measurements across time points (MTr-GBLUP) were jointly analyzed. Both formulations share the same linear model as follows:

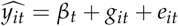

where 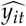 corresponds to the adjusted mean related to the i^th^ line evaluated for the t^th^ time point or trait, *β*_*t*_ is the fixed population mean effect for the t^th^ time point and/or trait, *git* is the genomic estimated breeding value (GEBV) of the i^th^ line evaluated for the t^th^ time point and/or trait, and *e*_*it*_ is the residual.

Considering the structure of multivariate GBLUP models with stacked trait and/or time subvectors like *g* = [*g*_**1**_, *g*_**2**_, … , *g*_*k*_]^*T*^ and *e* = [*e*_**1**_, *e*_**2**_, … , *e*_*k*_]^*T*^, in which *k* is the number of traits and/or time points, we assume that *g* ~ MVN(**0**, *G* ⊗ *A*) and 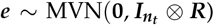, where *A* (*n*_*t*_ × *n*_*t*_) is the additive relationship matrix (VanRaden 2008) between *n*_*t*_ lines, ***G*** (*k* × *k*) and ***R*** (*k* × *k*) are unstructured variance-covariance matrices for genetic and residual effects, respectively. The ***A*** matrix was constructed using the 100,435 SNP markers with the A.mat function in the R package *rrBLUP 4.6* (Endelman 2011). Spectral decomposition was performed to transform the *A* matrix into positive definite. The procedure was based on the singular value decomposition of the *A* matrix, substitution of the negative values by a decreasing small constant (10^−4^), and reconstruction of the *A* matrix. Additional details on the spectral decomposition procedure are available in Calinński *et al.* (2005); dos Santos *et al.* (2016b). The MTi-GBLUP and MTr-GBLUP models were fitted using the R package *EMMREML 3.1* (Akdemir and Godfrey 2018).

### Cross-validation schemes

Two different cross-validation (CV) schemes were used to evaluate the predictive accuracy of the GP models. The first scheme used was stratified 5-fold CV for each individual trait (i.e., DBY or PH measured at a single time point). This procedure was based on stratifying the phenotypic and genotypic data of the lines into five non-overlapping folds, training the model with four folds (training set), and predicting the phenotypes of lines in the fold not included for training (test set) with only their genotypes as predictors in the trained model. This procedure was repeated until phenotypes from all five folds were predicted. Forward-chaining CV was used as a second scheme. In this scheme, data were split into time point subsets. The initial training set of five total was based on data from the first two time points (30 and 45 DAP), with the remaining time points (60, 75, 90, 105, and 120 DAP) comprising the test set. This procedure was repeated four more times to build new training sets until all time points except the last one (120 DAP) were included in the training set. The forward-chaining CV scheme was used to assess the accuracy of GP models to predict and identify the best set of lines for PH (tallest) prior to harvest.

The correlation (Pearson’s *r*) of adjusted means with predicted values was used to estimate predictive accuracy in both CV schemes. In the forward-chaining CV scheme, the predicted values were always obtained with the artificial bins effects from the previous time point used for training the DBN model. For the MTi-GBLUP and MTr-GBLUP models, the predicted values from the previous time point used for training were used as predicted values. For the BN and PBN models, which do not share information across time, the effects of artificial bins for each time point were used to compute the predicted values.

### Coincidence index

For the five GP models, coincidence indices (CIs) were constructed to evaluate the capacity for selecting the top 20% best performing lines for DBY when considering the rank of the PH adjusted mean values at each time point. The posterior values of the CI were calculated as the rate of successes between the top 20% best lines for PH at each time point and DBY. The CI was computed by assigning a ‘1’ to lines in the top 20% best lines for PH and DBY, or ‘0’ otherwise, then dividing the total number of successes (sum of ‘1’s) by the total number of lines at each posterior sample.

### Coincidence index based on lines

For the DBN model, we constructed a coincidence index based on lines (CIL) that used posterior samples from the adjusted mean values of PH across the seven developmental time points. This CIL was used to determine how early selection could be performed within-season to optimally reduce the length of the breeding cycle. Calculation of the CIL was based on the following steps: (i) identify the top 20% best lines for each posterior sample of the PH adjusted mean values at each time point; (ii) create for each posterior sample a one-hot vector encoding, assigning a ‘1’ to the best lines in the top 20% in the evaluated time point and at the end of the season (120 DAP), or ‘0’ otherwise; and (iii) compute the CIL as the total number of successes divided by the total number of posterior samples that each line appeared in the top 20% for the predicted (evaluated time points) and observed adjusted mean values (end of season).

### Data Availability

Genotypic and phenotypic data are available at CyVerse (http://datacommons.cyverse.org/browse/iplant/home/shared/GoreLab/dataFromPubs/dosSantos_BayesianNetworks_2019).

Scripts used in this study are available on GitHub (https://github.com/GoreLab/sorghum-multi-trait).

## RESULTS

### Phenotypic variation

We used a mixed linear model that accounted for the influence of environment and genotype-by-environment interaction to generate adjusted means for end-of-season DBY and PH measured at seven developmental time points over the growing season. Both DBY and the multiple PH measures had moderately high estimates of heritability on a line-mean basis (*h*^2^ > 0.48) across the four environments (Figure 2). The distribution of adjusted mean values for DBY was slightly skewed towards the left tail, with values centered on 23.55 t ha^−1^ (std=5.79). In comparison, the adjusted mean values for PH showed an expected growth pattern across the seven time points, with the population mean for PH changing from 25.6 (std=3.7) at 30 DAP to 350.7 cm (std=48.0) at 120 DAP. Indicative of an autoregressive trend of correlation over time, the weakest correlation (*r*) was observed between PH measures collected at 30 and 120 DAP (*r*=0.40), while 90 and 105 DAP were the two most strongly correlated time points (*r*=0.96). Correlations between DBY and PH varied from 0.10 to 0.31 across time points, suggesting an opportunity for recovering information across time points and/or between traits to improve the predictive accuracy of GP models.

**Figure 2.**
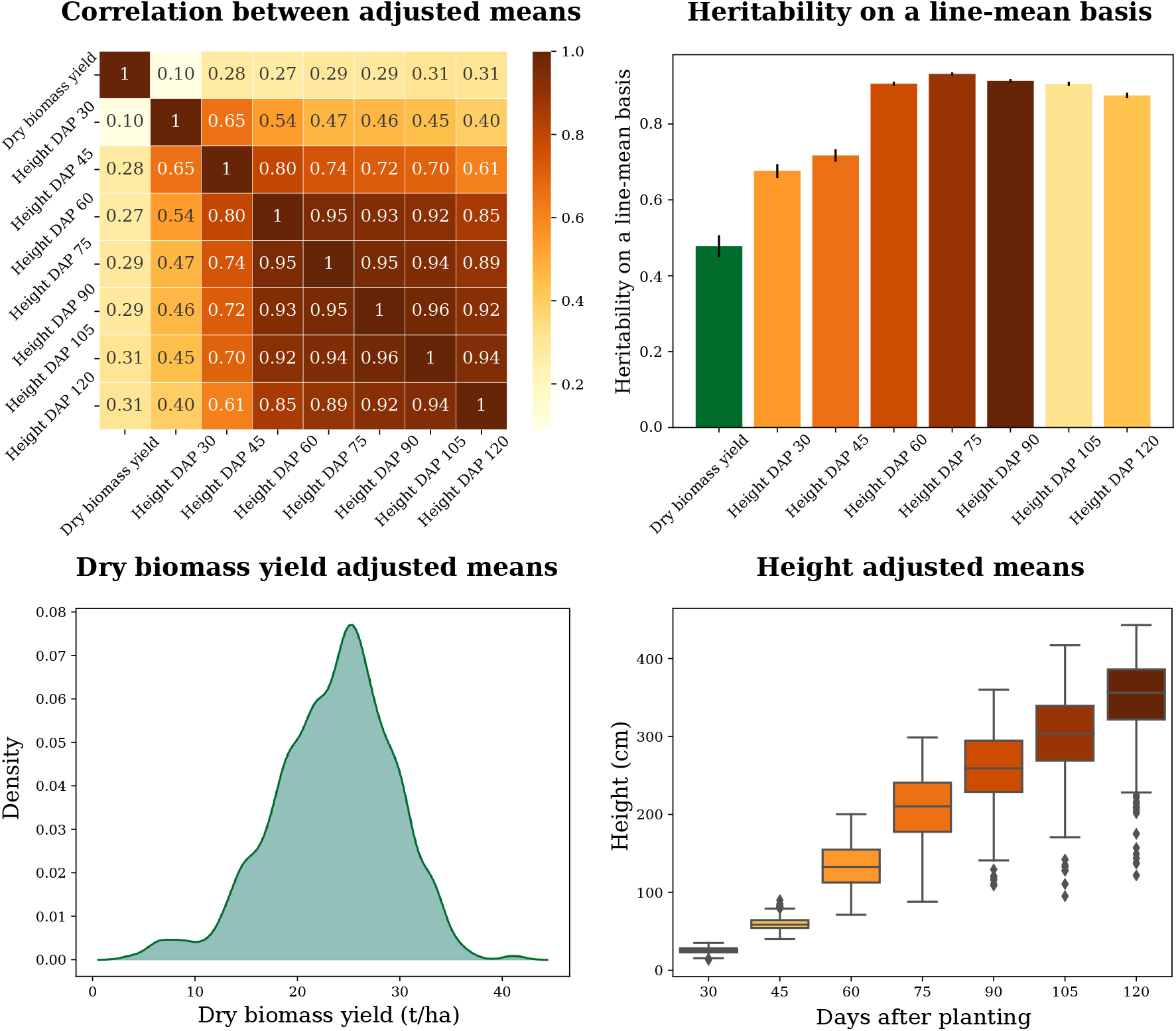
Summary statistics of evaluated phenotypes. Correlations between adjusted means, heritabilities, and distributions of adjusted means.

### Predictive accuracies from stratified 5-fold CV

We evaluated the accuracy of the BN, PBN, DBN, MTr-GBLUP, and MTi-GBLUP models for predicting DBY and PH measured throughout the growing season with a stratified 5-fold CV scheme. Prediction accuracies (0.48-0.51) of DBY were nearly identical for the BN, PBN, and MTr-GBLUP models (Table 1). When predicting PH at each of the seven developmental stages with the BN, PBN, MTi-GBLUP and MTr-GBLUP models, we found that accuracies gradually increased from 30 to 60 DAP, peaked at 60 DAP, and incrementally decreased from 60 to 120 DAP. Of these four models, MTi-GBLUP (0.56-0.74) and MTr-GBLUP (0.57-0.74) had prediction accuracies comparable to each other and slightly higher than those of BN (0.53-0.72) and PBN (0.52-0.72). Comparatively, the DBN model showed a randomly fluctuating trend of relatively slightly lower prediction accuracies for PH at all seven time points. The minor variation in predictive accuracy, especially after 45 DAP, suggests that early season prediction could be possible for PH.

**Table 1.**
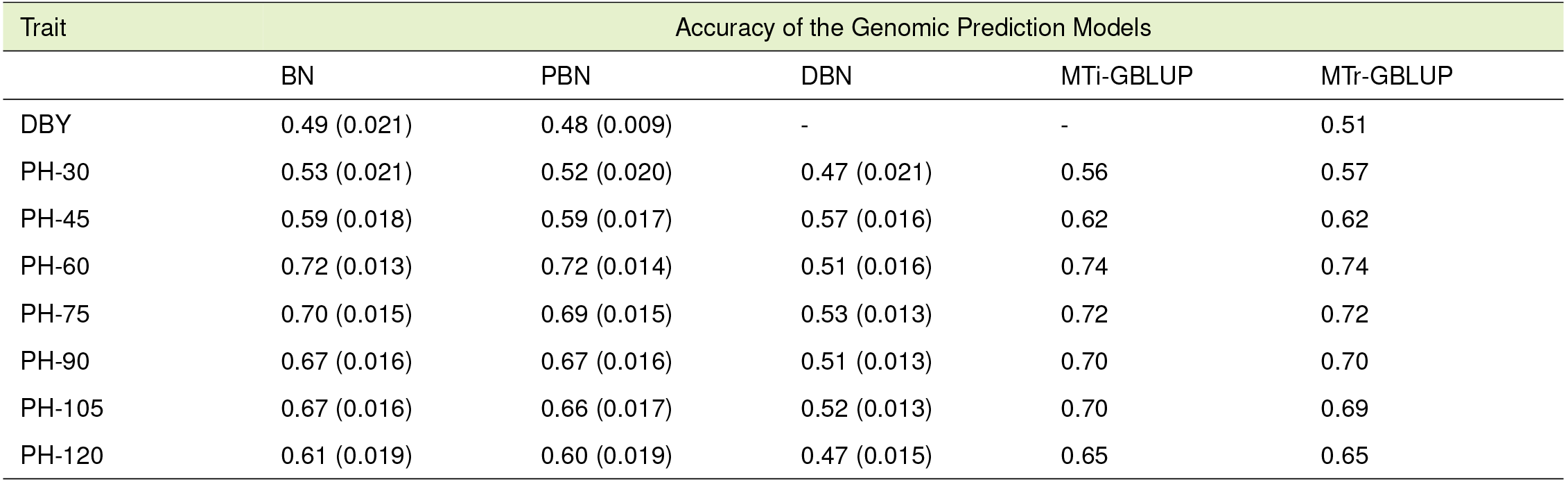
Prediction accuracies obtained from the 5-fold cross-validation scheme by training the Bayesian network (BN), pleiotropic Bayesian network (PBN), dynamic Bayesian network (DBN), multi time GBLUP (MTi-GBLUP) and multi trait GBLUP (MTr-GBLUP) models to predict dry biomass yield (DBY) collected at harvest and plant height (PH) measured across different days after planting (DAP). The standard deviation of the prediction accuracy obtained by each Bayesian model is reported within parentheses.

| Trait | Accuracy of the Genomic Prediction Models |  |  |  |  |
| --- | --- | --- | --- | --- | --- |
|  | BN | PBN | DBN | MTi-GBLUP | MTr-GBLUP |
| DBY | 0.49 (0.021) | 0.48 (0.009) | - | - | 0.51 |
| PH-30 | 0.53 (0.021) | 0.52 (0.020) | 0.47 (0.021) | 0.56 | 0.57 |
| PH-45 | 0.59 (0.018) | 0.59 (0.017) | 0.57 (0.016) | 0.62 | 0.62 |
| PH-60 | 0.72 (0.013) | 0.72 (0.014) | 0.51 (0.016) | 0.74 | 0.74 |
| PH-75 | 0.70 (0.015) | 0.69 (0.015) | 0.53 (0.013) | 0.72 | 0.72 |
| PH-90 | 0.67 (0.016) | 0.67 (0.016) | 0.51 (0.013) | 0.70 | 0.70 |
| PH-105 | 0.67 (0.016) | 0.66 (0.017) | 0.52 (0.013) | 0.70 | 0.69 |
| PH-120 | 0.61 (0.019) | 0.60 (0.019) | 0.47 (0.015) | 0.65 | 0.65 |

### Predictive accuracies from forward-chaining cross-validation

We performed a forward-chaining CV procedure to evaluate the accuracy of the five models to predict PH at unobserved time points. In general, the models showed high accuracy to predict the phenotypic values of the lines observed at the last time point (120 DAP) even when trained only with data from both 30 and 45 DAP (Figure 3). The BN and PBN models had similar prediction accuracies across all scenarios, ranging from 0.42 (BN, training: 45 DAP; predicting: 120 DAP) to 0.86 (PBN, training: 60 DAP; predicting: 75 DAP). In contrast, the DBN, MTi-GBLUP, and MTr-GBLUP models had substantially higher predictive accuracies compared to the BN and PBN models. The DBN model showed predictive accuracies varying from 0.6 (training slice: 30-45 DAP, predicting: 120 DAP) to 0.95 (training slice: 30-60 DAP; predicting: 75 DAP). The MTi-GBLUP prediction accuracies ranged from 0.63 (training slice: 30-45 DAP; predicting: 120 DAP) to 0.94 (training slice: 30-90 DAP; predicting: 105 DAP), and comparably, the MTr-GBLUP varied from 0.64 (training slice: 30-45 DAP; predicting: 120 DAP) to 0.94 (training slice: 30-90 DAP; predicting: 105 DAP). These results did not suggest any advantage for modelling dependence between PH and DBY in the PBN and MTi-GBLUP models; however, the results did suggest that the dependence between time points accounted for in the DBN, MTi-GBLUP and MTr-GBLUP models improved the prediction accuracy of PH.

**Figure 3.**
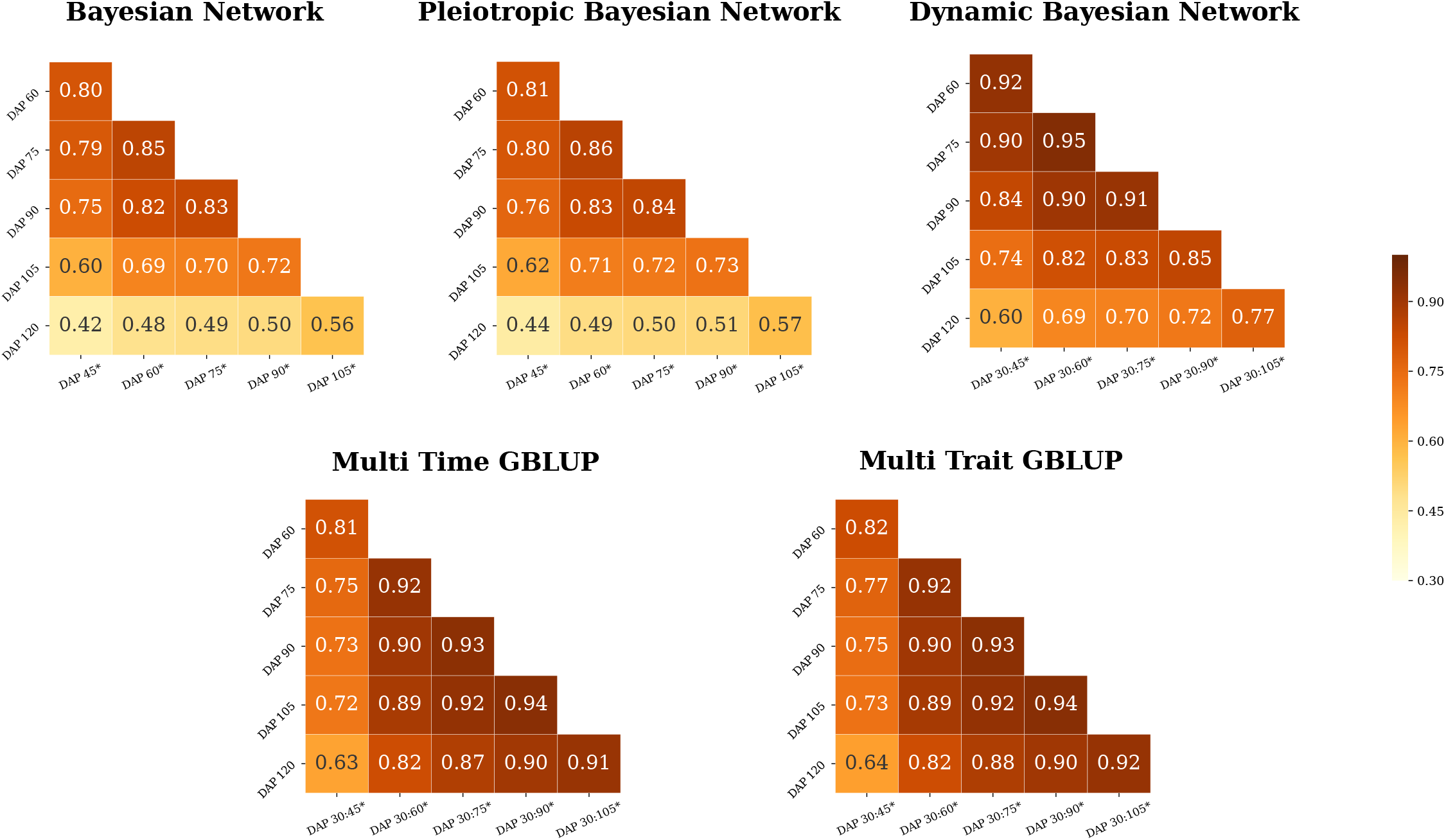
Genomic prediction of plant height with forward chaining cross-validation. Prediction accuracies were obtained from prediction models exploiting single (Bayesian Network and Pleiotropic Bayesian Network) or multiple time points (Dynamic Bayesian Network, Multi Time GBLUP, and Multi Trait GBLUP). The horizontal axis represents the slice (:) of the time interval used for training the models with multiple time points and the vertical axis the testing data. The ‘*’ symbol denotes the days after planting (DAP) time point used to obtain the adjusted means.

### Coincidence indexes

The high predictive performance of the DBN, MTri-GBLUP, and MTr-GBLUP models, which exploited multiple PH measurements over initial growth stages, incentivized us to investigate how the rank of the lines varied across the different time points. To that end, we evaluated how PH measures over time (secondary traits) could be informative for performing indirect selection of DBY (target trait) through the calculation of coincidence indices (CIs). The posterior distribution of the CIs showed an overlapping pattern over time for the three Bayesian models (Figure 4), with most of them ranging from 0.18 to 0.34. This implied that the ranking of the lines for PH did not change significantly from early to late growth stages. Additionally, the MTi-GBLUP CI (first-level target trait: DBY; second-level target trait: PH) ranged from 0.25 (training slice: 30-105 DAP) to 0.27 (training slice: 30-45 DAP), and MTr-GBLUP varied from 0.26 (training slice: 30-105 DAP) to 0.28 (training slice: 30-45 DAP). These results suggest that the initial growth stage ranging from 30 to 45 DAP could be an optimal stage of development for the early selection of PH and indirect selection of DBY based on the ordering of the PH adjusted means over the growing season.

**Figure 4.**
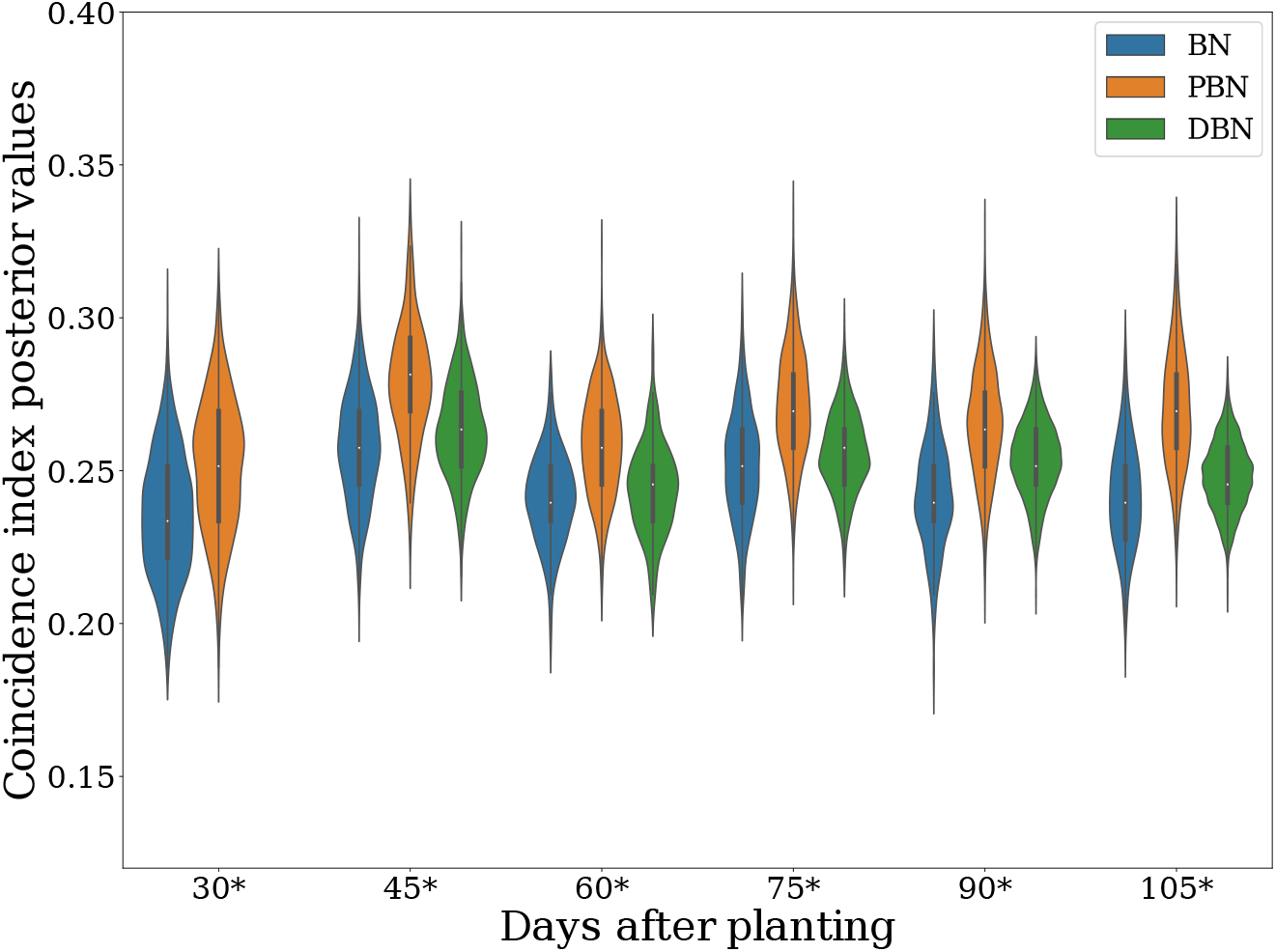
The calculated coincidence index (CI) at multiple developmental stages. The CILs for selecting the top 20% for dry biomass yield at specific developmental stages were calculated using as reference the adjusted mean values obtained by training the Bayesian network (BN), pleiotropic Bayesian network (PBN), and dynamic Bayesian network (DBN) models with the plant height time series. The CIs from the multi time GBLUP (MTi-GBLUP) and multi trait GBLUP (MTr-GBLUP) models were not plotted (point estimates reported in the text). The ‘*’ symbol denotes the time point estimates used to obtain the adjusted means as expected values for indirect selection. For the DBN model that leveraged multiple time points, the symbol * denotes the last time point used for training with the earlier time points also considered in the model.

To gain more insight and empirical evidence to support the hypothesis of early selection for PH with measures from 30 and 45 DAP, we developed a coincidence index based on lines (CIL) using the posterior values from the DBN model that achieved optimal performance among the Bayesian models tested in the forward-chaining CV. The CIL allowed us to better understand phenotypic plasticity through assessing how the expected rank of the lines at the end of the season agreed with their ranking at earlier growth stages. The closer that the CIL is to one, the more likely the line is expected to be at the top 20% for PH at the end of the season. We plotted lines with CIL > 0.5, fixed their ordering (training slice: 30-45 DAP), and displayed the CILs from other time slices in the same order (Figure 5). The CILs showed the expected trend of increasing the chance of lines to be in the top 20% after 45 DAP, which indicated that the ranking of the top lines had not majorly changed over time.

**Figure 5.**
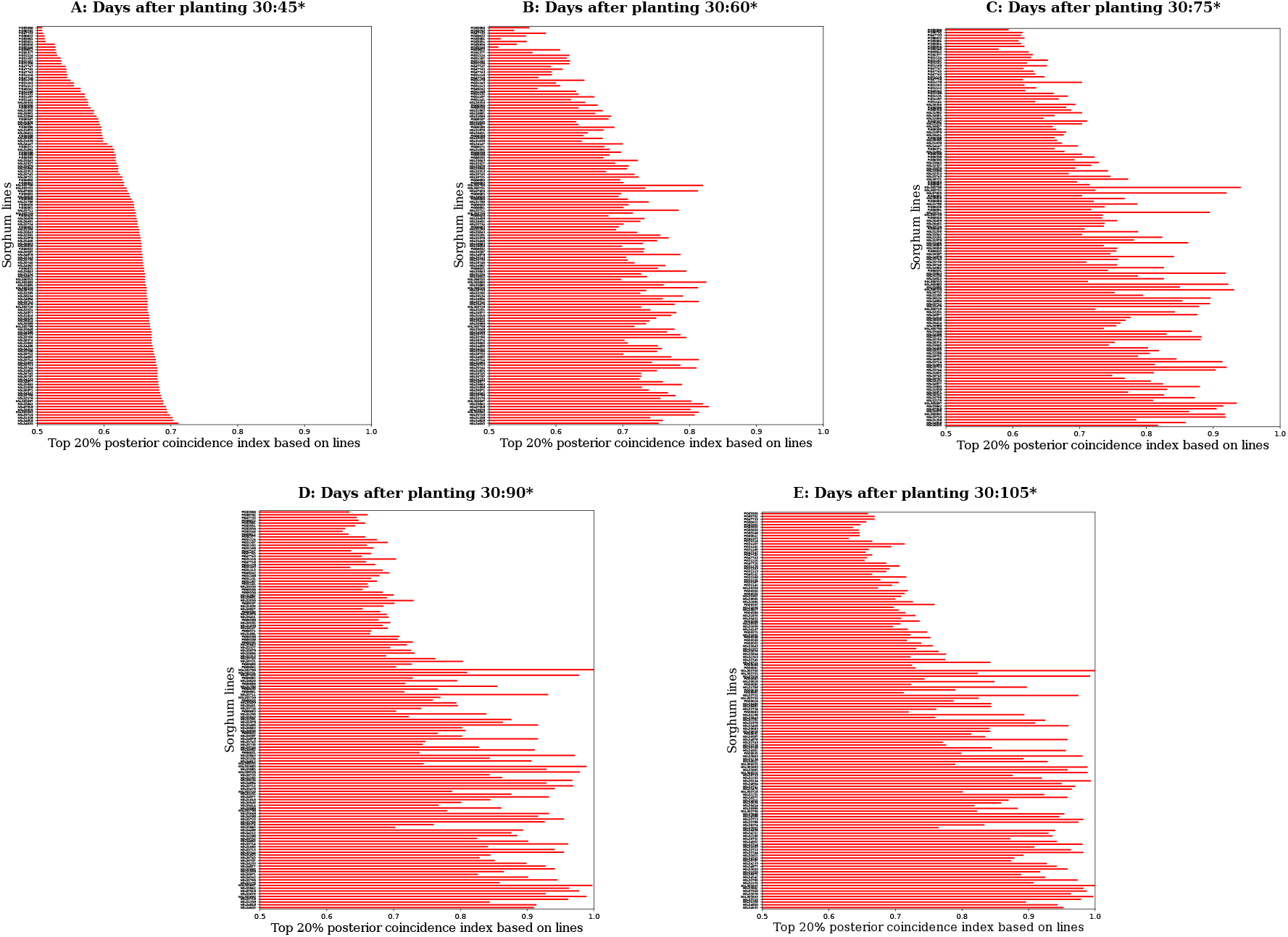
The calculated coincidence index based on lines (CIL) at multiple developmental stages. The top 20% posterior CILs from the results of the dynamic Bayesian network (DBN) are shown. The rank order of the lines in subplot A was fixed for subplots B, C, D, and E to understand phenotypic plasticity over time. Only lines with a CIL>0.5 were plotted. The ‘*’ symbol denotes the days after planting (DAP) time point that was used to obtain the adjusted means.

## DISCUSSION

Biomass sorghum is a promising bioenergy feedstock because of its extensive genetic diversity, high biomass yield potential, and strong tolerance to environmental stress. Sorghum is evolutionarily related to key bioenergy grasses, including maize, sugarcane, switchgrass, and *Miscanthus* spp., making it a potentially important diploid model to inform the genetic improvement of these other bioenergy crops (Morris *et al.* 2013; Brenton *et al.* 2016). Despite sorghum’s appealing features as both a crop and model species, few studies have focused on genetically modelling its growth patterns and leveraging this information for breeding optimization. In this study, we investigated a diverse panel of 839 sorghum lines genotyped with 100,435 SNP markers that was evaluated for a PH time series and DBY in four environments. With these collected data, we evaluated several GP models for exploiting genetic information over time and/or between correlated traits to improve prediction accuracies compared to models that assumed independence. Our implemented Bayesian models allowed us to estimate the level of uncertainty in determining optimal time points when developing breeding strategies for early selection of PH within season, as well as the indirect selection of DBY in combination with the repeated measures of PH as secondary traits.

To conduct GP of DBY and PH, we used both PGM and multivariate mixed linear model approaches to better model growth dynamics (Bishop 2013; Henderson and Quaas 1976; dos Santos *et al.* 2016b). Due to the high computational cost of the PGM approach, we modified the artificial bins method of Xu (2013) to reduce the dimensionality of the SNP marker matrix through a PCA. This modified approach reduced the number of parameters needed to train the different PGM architectures by a 100-fold. Also, this procedure in other scenarios has the flexibility for predictions even when the number of loci pooled is different between the training and testing sets. Indicative of minimal information loss, there were negligible differences in prediction accuracies achieved by the MTi-GBLUP and MTr-GBLUP models that used the 100,000 SNP markers to compute the relationship matrix compared to those of the BN and PBN models reliant on the 1,000 artificial bins. Also, the artificial bins approach did not compromise the results from the DBN model, as indicated by the similarity of its obtained prediction accuracies with those from the multivariate GBLUP models in the forward-chaining CV scheme.

We initiated a model-based machine learning approach for GP analysis by first defining the baseline of PGMs (Bishop 2013). Several studies have shown a similar level of predictive performance between PGMs that assume either a common or specific normal prior for each marker effect (de Los Campos *et al.* 2013; Heslot *et al.* 2015; Ferrão *et al.* 2018). Therefore, we parsimoniously used a common normal prior for the effects of all artificial bins, resulting in a BN model that had less unknown parameters but a slower MCMC process to obtain posterior draws. The BN model can be considered a non-conjugate form of the Bayesian linear regression model (de Los Campos *et al.* 2013) that automatically learns the hyperparameters of priors from the data. This model also has Cauchy priors truncated to the 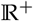 on the scale components, which avoids sampling implausible standard deviation values (Gelman *et al.* 2014). Although the main disadvantage of the BN model formulation is that its architecture cannot recover information between traits or time points, the BN model can be useful for analyzing highly unbalanced data as is frequently observed for large-scale field trials in the plant breeding industry.

To improve the performance of the BN model, we novelly developed the PBN model by connecting two representations of the BN model with hidden variables (nodes in the graph representing artificial bins effects influencing both traits), allowing a conditional relationship between the likelihood functions of PH and DBY to be established. Despite our efforts to estimate pleiotropic effects with the PBN model, the interpretation of the effects as pleiotropic should be carefully interpreted, especially due to the challenge of differentiating between pleiotropy and tight linkage. Gianola *et al.* (2015) theorized that the linkage disequilbrium (LD) between markers and quantitative trait loci (QTL), LD between QTL controlling different traits, or LD between markers linked to QTL controlling different traits can make it difficult to partition pleiotropic effects in GP models. The presence of such complex LD structure could explain the minor difference in predictive accuracy between the BN and PBN models when attempting to partition genetics effects influencing single versus multiple traits. This interpretation is supported by the finding that prediction accuracies obtained by the MTi-GBLUP (PH time series) and MTr-GBLUP (DBY and PH time series) models were similar to each other. Additionally, the lower than expected performance of the PBN model relative to the BN model could be attributed to the modest correlations observed between DBY and the multiple PH traits (*r* = 0.10 - 0.31). Simulation studies have shown that genetic correlations weaker than 0.5 do not provide marked improvements to the prediction accuracy of GP models used for multiple traits (Calus and Veerkamp 2011; dos Santos *et al.* 2016b). Further studies analyzing traits having strong genetic correlations with DBY are needed to better understand how to further improve the prediction accuracy of the PBN model.

The DBN model, a variant of hidden Markov models that is intended for modeling continuous variables (Murphy 2013), was used to exploit the effects of artificial bins from the prediction of PH at earlier time points. This connection of genetic information between time points was crucial for dramatically improving the performance of the Bayesian framework in the forward-chaining CV scheme. Moreover, the predictions using the posterior mean of the DBN model allowed us to obtain predictions as precise as the point estimates from the MTi-GBLUP model and construct indices with posterior samples to identify optimal time points for indirect selection. The strong correlation between multiple time points for PH is quite possibly the main factor that favored the improvement of prediction accuracy for the DBN model compared to the BN and PBN models that did not leverage information over time. In contrast to the high predictive performance achieved in the forward-chaining CV scheme, relatively lower predictive accuracies were observed for the DBN model in the 5-fold CV scheme, especially when removing lines across all time points to split the data into training and testing sets. This is because the splitting did not allow the DBN model to learn with precision the effects of the artificial bins between time points and added noise to the artificial bin estimates. Despite the sharing of genetic signals over time, these findings suggest that the DBN model should not be used when lines are completely unobserved across all time points (*Burgueño et al.* 2012; Dias *et al.* 2018; Fernandes *et al.* 2018). This reduced level of prediction performance because of unobserved lines has also been previously reported for multivariate GP models tested with a 5-fold CV scheme (Burgueño *et al.* 2012; Dias *et al.* 2018; Fernandes *et al.* 2018).

Even though modeling growth dynamics is an important area of research, there have been a limited number of studies in plants that have analyzed longitudinal phenotypes related to growth rate with GP models. In a greenhouse study of 357 diverse rice (*Oryza sativa* L.) accessions, Campbell *et al.* (2018) developed GP models with a first- or second-order Legendre polynomial to predict the sum of “plant pixels” from image-based phenotyping as a daily estimate of shoot biomass during initial growth stages (13 to 33 days after transplant), resulting in an 11.6% improvement in prediction accuracies relative to a single time point analysis. Despite the advantage of fitting a nonlinear random regression model with phenotypic data that showed an exponential curve over 20 days of early vegetative growth, this procedure is limited to only early developmental stage phenotypes such as shoot biomass that follow an exponential growth curve. In our study, PH was measured throughout the entire growing season without a focus on the early vegetative stage, thus the collected PH data did not have an exponential shape. The modelling of genetic effects as either a linear additive function over time or though the *G* var-cov matrix might be the main factor causing the 36.4-52.4% (training slice: 30-45 DAP; predicting: 120 DAP) improvements in prediction accuracy of the DBN, MTi-GBLUP, and MTr-GBLUP models relative to the BN and PBN models. When analyzing repeated measures of height collected from an interior spruce (*Picea engelmannii* x *glauca*) population of 769 trees at six sparse time points over a period of 37 years, Ratcliffe *et al.* (2015) observed minor differences in prediction performance among the evaluated BayesC*π*, ridge regression (rrBLUP), and generalized ridge regression models. These findings are contrary to our evaluated PGMs that recovered genetic information between time points that showed substantial improvement compared to the BN model—a Bayesian formulation of the rrBLUP model.

The implemented PGMs provided a powerful modelling framework to infer uncertainty based on well-established probability theory (Murphy 2013; Bishop 2013), allowing us to define optimal time points for early within-season selection. To this end, the posterior distribution of the CIs was used to show that the rank order of the sorghum lines changed minimally after 45 DAP, which suggested an opportunity to indirectly select for DBY based on early season PH measures as secondary traits. The Bayesian CIs allowed the overlapping pattern of posterior values to be assessed for identifying optimal time points for selection contrary to the point estimate approach of Hamblin and Zimmermann (1986) applied to genomic prediction of a PH time series and DBY in sorghum by Fernandes *et al.* (2018). The Bayesian CIs also did not require resampling to build the index. In addition, we used the DBN model to evaluate the phenotypic plasticity of PH with the CILs at the population level. The pattern of phenotypic plasticity shown by the CILs confirmed the general findings of the CIs—the ranking of lines by PH changed minimally from early (45 DAP) to late (120 DAP) time points. Early within-season selection for PH could help to efficiently accelerate the breeding process. Considering the Breeder’s equation ΔG = i h *σ*_*A*_/L (Li *et al.* 2018), reducing the length of the generation (L) for given a selection intensity (i) could provide a new avenue for increasing genetic gain per unit of time through early indirect selection of PH and DBY at 45 DAP in these tested environments.

Given that the order of lines ranked by PH minimally changed across the measured plant developmental stages and had a moderate coincidence with rankings based on DBY, we propose a two-level indirect selection framework: (i) fit the DBN model, which of the Bayesian models had the best performance for predicting future measures, to learn posterior values of the GEBVs for PH repeated measures in initial growth stages and obtain a precise estimate of the ranking of each line at the end of the season; (ii) compute CIs and CILs to evaluate the extent to which the rank order of lines change; (iii) use GEBVs of the last time point used for training (e.g., 45 DAP for our tested environment) as a secondary trait; (iv) perform indirect selection for PH at the end of the season as a first-level target trait; and (v) perform indirect selection for DBY as the second-level target trait. This selection approach together with the trait-assisted genomic selection approach proposed by Fernandes *et al.* (2018) may allow the end-of-season rank order of observed and eventually unobserved lines to be accurately predicted early in the growing season. When integrated with high-throughput phenotyping platforms, the two-level indirect selection framework has the potential to further accelerate selection cycles and support a larger number of evaluated families. This could be accomplished by deploying low-cost ground rovers (e.g., Earthsense, https://www.earthsense.co) for the early season measurement of height and other traits genetically correlated with DBY on field-grown plants in combination with off-season winter nurseries and greenhouses with automated phenotyping systems (e.g., Lemnatec, http://www.lemnatec.com; Photon Systems Instruments, http://www.psi.cz) and optimized LED lights for speed breeding (Watson *et al.* 2018).

## CONCLUSION

We analyzed phenotypic measures over time for PH and DBY at the end of the season to design a novel indirect selection scheme. To that end, we developed and evaluated novel Bayesian networks for GP that were used to better model and understand phenotypic plasticity of PH across different plant developmental stages. The GP models showed minor differences in prediction accuracies for the 5-fold CV scheme. In stark contrast, in the forward-chaining CV scheme, we observed a 36.4-52.4% improvement in prediction accuracy of the DBN and multivariate GBLUP models (train on 30-45 DAP, predict 120 DAP) compared to the BN and PBN models that assumed independence over time. The Bayesian models were used to show that the ranking of lines changed minimally after 45 DAP based on the CI and CIL, serving as novel approaches to understand ranking dynamics with repeated measures. These results suggest that in these environments 45 DAP is an optimal developmental stage for imposing a two-level indirect selection framework for biomass sorghum. Such that indirect selection for end of season PH (first-level target trait) and DBY (second-level target trait) could be performed based on the ranking of lines by PH at 45 DAP (secondary trait).

## ACKNOWLEDGEMENTS

This work was supported by FAPESP (Fundação de Amparo à Pesquisa do Estado de São Paulo) Grant 2017/03625-2 and 2017/25674-5. CAPES (Coordenação de Aperfeiçoamento de Pessoal de Nível Superior) Finance Code 001. CNPq (Conselho Nacional de Desenvolvimento Científico e Tecnológico). The information, data, or work presented herein was funded in part by the Advanced Research Projects Agency-Energy (ARPA-E), U.S. Department of Energy, under Award Number DE-AR0000598. This work was also supported by the USDA-ARS. The views and opinions of authors expressed herein do not necessarily state or reflect those of the United States Government or any agency thereof.

## AUTHOR CONTRIBUTIONS

J.P.R.D.S. and M.A.G. co-wrote the manuscript; J.P.R.D.S. led data analysis; P.J.B. and S.B.F. designed and managed field trials; P.J.B. collected sequencing and phenotyping data; S.B.F. collected phenotypic data; R.L. analyzed sequencing data; J.P.R.D.S. developed genomic prediction models, codes, and statistical analysis methods; E.S.B. and A.A.F.G. consulted on data analysis and provided support in the development of genomic prediction models and project coordination; M.A.G. overall project management, design, coordination, oversaw data analysis; all authors contributed to the critical review of the manuscript.

## LITERATURE CITED

Akdemir, D. and O. U. Godfrey, 2018 EMMREML: Fitting Mixed Models with Known Covariance Structures (Version 3.1).

Bae, H., S. Monti, M. Montano, M. H. Steinberg, T. T. Perls, et al., 2016 Learning Bayesian Networks from Correlated Data. Scientific reports 6: 1–14.

Bao, Y., L. Tang, M. W. Breitzman, M. G. Salas Fernandez, and P. S. Schnable, 2019 Field-based robotic phenotyping of sorghum plant architecture using stereo vision. Journal of Field Robotics 36: 397–415.

Bishop, C. M., 2013 Model-based machine learning. Phil Trans R Soc A 371: 1–17.

Brenton, Z. W., E. A. Cooper, M. T. Myers, R. E. Boyles, N. Shakoor, et al., 2016 A genomic resource for the development, improvement, and exploitation of sorghum for bioenergy. Genetics 204: 21–33.

Browning, B. and S. Browning, 2016 Genotype imputation with millions of reference samples. The American Journal of Human Genetics 98: 116 – 126.

Buckler, E. S., D. C. Ilut, X. Wang, T. Kretzschmar, M. A. Gore, et al., 2016 rampseq: Using repetitive sequences for robust genotyping. bioRxiv.

Burgueño, J., G. de los Campos, K. Weigel, and J. Crossa, 2012 Genomic Prediction of Breeding Values when Modeling Genotype × Environment Interaction using Pedigree and Dense Molecular Markers. Crop Science.

Butler, D. G., B. R. Cullis, A. R. Gilmour, and B. J. Gogel, 2009 ASReml-R reference manual.

Caliński, T., S. Czajka, Z. Kaczmarek, P. Krajewski, and W. Pilarczyk, 2005 Analyzing multi-environment variety trials using randomization-derived mixed models. Biometrics 61: 448–455.

Calus, M. P. and R. F. Veerkamp, 2011 Accuracy of multi-trait genomic selection using different methods. Genetics Selection Evolution 43: 26.

Campbell, M., H. Walia, and G. Morota, 2018 Utilizing random regression models for genomic prediction of a longitudinal trait derived from high-throughput phenotyping. Plant Direct 2: e00080.

Carpenter, B., A. Gelman, M. Hoffman, D. Lee, B. Goodrich, et al., 2017 Stan: A probabilistic programming language. Journal of Statistical Software, Articles 76: 1–32.

Davey, J. W., P. A. Hohenlohe, P. D. Etter, J. Q. Boone, J. M. Catchen, et al., 2011 Genome-wide genetic marker discovery and genotyping using next-generation sequencing. Nature Reviews Genetics 12: 499–510.

de Los Campos, G., J. M. Hickey, R. Pong-Wong, H. D. Daetwyler, and M. P. L. Calus, 2013 Whole-genome regression and prediction methods applied to plant and animal breeding. Genetics 193: 327–345.

Dias, K. O. G., S. A. Gezan, C. T. Guimarães, A. Nazarian, L. da Costa e Silva, et al., 2018 Improving accuracies of genomic predictions for drought tolerance in maize by joint modeling of additive and dominance effects in multi-environment trials. Heredity.

dos Santos, J. P., L. Paulo, M. Pires, R. Coelho, and D. C. Vasconcellos, 2016a Genomic selection to resistance to Stenocarpella maydis in maize lines using DArTseq markers. BMC Genetics 17: 1–10.

dos Santos, J. P. R., R. C. De Castro Vasconcellos, L. P. M. Pires, L. Balestre, and R. G. Von Pinho, 2016b Inclusion of dominance effects in the multivariate GBLUP model. PLoS ONE 11: 1–21.

Elshire, R. J., J. C. Glaubitz, Q. Sun, J. A. Poland, K. Kawamoto, et al., 2011 A Robust, Simple Genotyping-by-Sequencing (GBS) Approach for High Diversity Species. PLoS ONE 6: e19379.

Endelman, J. B., 2011 Ridge regression and other kernels for genomic selection with r package rrblup. Plant Genome 4: 250–255.

Fernandes, S. B., K. O. G. Dias, D. F. Ferreira, and P. J. Brown, 2018 Efficiency of multi-trait, indirect, and trait-assisted genomic selection for improvement of biomass sorghum. Theoretical and Applied Genetics 131: 747–755.

Ferrão, L. F. V., R. G. Ferrão, M. A. G. Ferrão, A. Fonseca, P. Carbonetto, et al., 2018 Accurate genomic prediction of Coffea canephora in multiple environments using whole-genome statistical models. Heredity.

Foley, J. a., N. Ramankutty, K. a. Brauman, E. S. Cassidy, J. S. Gerber, et al., 2011 Solutions for a cultivated planet. Nature 478: 337–342.

Garcia, A. a. F., M. Mollinari, T. G. Marconi, O. R. Serang, R. R. Silva, et al., 2013 SNP genotyping allows an in-depth characterization of the genome of sugarcane and other complex autopolyploids. Scientific Reports 3: 3399.

Gelman, A., J. B. Carlin, H. S. Stern, and D. B. Rubin, 2014 Bayesian Data Analysis.

Gianola, D., 2013 Priors in whole-genome regression: the bayesian alphabet returns. Genetics 194: 573–96.

Gianola, D., G. de los Campos, W. G. Hill, E. Manfredi, and R. Fernando, 2009 Additive genetic variability and the Bayesian alphabet. Genetics 183: 347–363.

Gianola, D., G. de los Campos, M. A. Toro, H. Naya, C.-C. Schön, et al., 2015 Do molecular markers inform about pleiotropy? Genetics 201: 23–29.

Glaubitz, J. C., T. M. Casstevens, F. Lu, J. Harriman, R. J. Elshire, et al., 2014 Tassel-gbs: A high capacity genotyping by sequencing analysis pipeline. PLOS ONE 9: 1–11.

Goodfellow, I., Y. Bengio, and A. Courville, 2016 Deep Learning. MIT Press, http://www.deeplearningbook.org.

Hamblin, J. and M. d. O. Zimmermann, 1986 Breeding common bean for yield in mixtures. Plant Breeding Reviews 4: 245–272.

Hamelryck, T., 2012 Bayesian Methods in Structural Bioinformatics.

Han, B., X.-w. Chen, Z. Talebizadeh, and H. Xu, 2012 Genetic studies of complex human diseases: characterizing SNP-disease associations using Bayesian networks. BMC Systems Biology 6: 1–12.

Henderson, C. R. and R. L. Quaas, 1976 Multiple trait evaluation using relatives’ records. Journal of Animal Science 43: 1188–1197.

Heslot, N., J. L. Jannink, and M. E. Sorrels, 2015 Perspectives for Genomic Selection Applications and Research in Plants. Crop Science 55: 1–12.

Hoffman, M. D. and A. Gelman, 2014 The No-U-Turn Sampler: Adaptively Setting Path Lengths in Hamiltonian Monte Carlo. Journal of Machine Learning Research 15: 1593–1623.

Holland, J., W. Nyquist, and C. Cervantes, 2003 Estimating and interpreting heritability for plant breeding: An update. plant breeding reviews vol. 22. Technical report.

Hung, H., C. Browne, K. Guill, N. Coles, M. Eller, et al., 2012 The relationship between parental genetic or phenotypic divergence and progeny variation in the maize nested association mapping population. Heredity 108: 490.

Jia, Y. and J.-L. Jannink, 2012 Multiple-trait genomic selection methods increase genetic value prediction accuracy. Genetics 192: 1513–1522.

Langmead, B. and S. L. Salzberg, 2012 Fast gapped-read alignment with bowtie 2. Nature methods 9: 357.

Lawrence, C. J. and V. Walbot, 2007 Translational Genomics for Bioenergy Production from Fuelstock Grasses: Maize as the Model Species. The Plant Cell 19: 2091–2094.

Li, H., A. Rasheed, L. T. Hickey, and Z. He, 2018 Fast-forwarding genetic gain. Trends in Plant Science 23: 184 – 186.

Loman, N. J., J. Quick, and J. T. Simpson, 2015 A complete bacterial genome assembled de novo using only nanopore sequencing data. bioRxiv 12: 015552.

Lynch, M., B. Walsh, et al., 1998 Genetics and analysis of quantitative traits, volume 1. Sinauer Sunderland, MA.

Mace, E. S., S. Tai, E. K. Gilding, Y. Li, P. J. Prentis, et al., 2013 Whole-genome sequencing reveals untapped genetic potential in Africa’s indigenous cereal crop sorghum. Nature Communications 4: 1–9.

Meuwissen, T. H. E., B. J. Hayes, and M. E. Goddard, 2001 Prediction of total genetic value using genome-wide dense marker maps. Genetics 157: 1819–1829.

Morris, G. P., P. Ramu, S. P. Deshpande, C. T. Hash, T. Shah, et al., 2013 Population genomic and genome-wide association studies of agroclimatic traits in sorghum. Proceedings of the National Academy of Sciences of the United States of America 110: 453–458.

Mullet, J., D. Morishige, R. McCormick, S. Truong, J. Hilley, et al., 2014 Energy sorghum–a genetic model for the design of C4 grass bioenergy crops. Journal of Experimental Botany 65: 3479–89.

Muraya, M. M., J. Chu, Y. Zhao, A. Junker, C. Klukas, et al., 2017 Genetic variation of growth dynamics in maize (zea mays l.) revealed through automated non-invasive phenotyping. The Plant Journal 89: 366–380.

Murphy, K. P., 2013 Machine learning : a probabilistic perspective. MIT Press, Cambridge, Mass. [u.a.].

Neapolitan, R., D. Xue, and X. Jiang, 2013 Modeling the altered expression levels of genes on signaling pathways in tumors as causal bayesian networks. Cancer Informatics 13: 77–84.

Okeke, U. G., D. Akdemir, I. Rabbi, P. Kulakow, and J.-L. Jannink, 2017 Accuracies of univariate and multivariate genomic prediction models in african cassava. Genetics Selection Evolution 49: 88.

Patterson, H. D. and R. Thompson, 1971 Recovery of inter-block information when block sizes are unequal. Biometrika 58: 545–554.

Pauli, D., S. C. Chapman, R. Bart, C. N. Topp, C. J. Lawrence-Dill, et al., 2016 The quest for understanding phenotypic variation via integrated approaches in the field environment. Plant Physiology 172: 622–634.

Ratcliffe, B., O. G. El-Dien, J. Klapste, I. Porth, C. Chen, et al., 2015 A comparison of genomic selection models across time in interior spruce (Picea engelmannii × glauca) using unordered SNP imputation methods. Heredity 115: 547–555.

Serang, O., M. Mollinari, and A. A. F. Garcia, 2012 Efficient exact maximum a posteriori computation for Bayesian SNP genotyping in polyploids. PLoS ONE 7: 1–13.

Su, C., A. Andrew, M. R. Karagas, and M. E. Borsuk, 2013 Using Bayesian networks to discover relations between genes, environment, and disease. BioData Mining 6: 1–21.

Team, S. D., 2018 PyStan: the Python interface to Stan, Version 2.17.1.0..

Valluru, R., E. E. Gazave, S. B. Fernandes, J. N. Ferguson, R. Lozano, et al., 2018 Leveraging mutational burden for complex trait prediction in sorghum. bioRxiv.

VanRaden, P., 2008 Efficient methods to compute genomic predictions. Journal of Dairy Science 91: 4414 – 4423.

Vermerris, W., 2011 Survey of Genomics Approaches to Improve Bioenergy Traits in Maize, Sorghum and Sugarcane. Journal of Integrative Plant Biology 53: 105–119.

Watson, A., S. Ghosh, M. J. Williams, W. S. Cuddy, J. Simmonds, et al., 2018 Speed breeding is a powerful tool to accelerate crop research and breeding. Nature plants 4: 23.

Wolak, M. E., 2012 nadiv: an R package to create relatedness matrices for estimating non-additive genetic variances in animal models. Methods in Ecology and Evolution 3: 792–796.

Xu, S., 2013 Genetic mapping and genomic selection using recombination breakpoint data. Genetics 195: 1103–1115.

Yu, X., X. Li, T. Guo, C. Zhu, Y. Wu, et al., 2016 Genomic prediction contributing to a promising global strategy to turbocharge gene banks. Nature Plants 2: 1–7.

